# retroLEAP: rAAV-based retrograde trans-synaptic labeling, expression and perturbation

**DOI:** 10.1101/537928

**Authors:** Janine K Reinert, Ivo Sonntag, Hannah Sonntag, Rolf Sprengel, Patric Pelzer, Sascha Lessle, Michaela Kaiser, Thomas Kuner

**Author notes:** Correspondence: Janine K Reinert, Ivo Sonntag. These authors contributed equally to this work (Janine K Reinert, Ivo Sonntag).

## Abstract

Identifying and manipulating synaptically connected neurons across brain regions remains a core challenge in understanding complex nervous systems. RetroLEAP is a novel approach for <u>retro</u>grade trans-synaptic <u>L</u>abelling, <u>E</u>xpression <u>A</u>nd <u>P</u>erturbation. GFP-dependent recombinase (Flp-DOG) detects trans-synaptic transfer of GFP-tetanus toxin heavy chain fusion protein (GFP-TTC) and activates expression of any gene of interest in synaptically connected cells. RetroLEAP overcomes existing limitations, is non-toxic, highly flexible, efficient, sensitive and easy to implement.

## Main

To understand complex neuronal circuits, it is essential to precisely identify and manipulate neurons forming synaptically connected networks at different scales, ranging from local to brain-wide circuits. Traditionally, tracing approaches distinguish anterograde and retrograde tracers. The former refers to labelling via projection targets of neuronal axons and the latter to labelling via synaptic inputs connected to a dendritic tree^1^. Anterograde tracing makes use of fluorescent molecules or genetically expressed fluorophores to visualize axons and hence identify projections to target areas across the nervous system^2^. However, these tracers only allow for the identification of putative input areas, as they do not cross synaptic connections. In contrast, retrograde tracing refers to the labelling of the neuron via its axon, while retrograde trans-synaptic labelling specifically refers to the transfer of the label from the postsynaptic neuron expressing the label (donor cell) to connected presynaptic neurons (acceptor cell)^3^.

Current approaches for retrograde trans-synaptic tracing can be classified into three categories, each based on a different principle. Firstly, tracer proteins like wheat germ agglutinin (WGA) or the non-toxic heavy chain of the tetanus toxin (TTC) that are able to cross synapses between donor and acceptor cells in a retrograde manner^4,5^. Secondly, recombinant viruses that cause a secondary infection of the acceptor cell by retrogradely crossing a synapse like herpes simplex virus (HSV), rabies virus (RV) or vesicular stomatitis virus (VSV)^1,6,7^. Thirdly, axon terminals can take up cholera toxin subunit B (CtB) or get directly infected with canine associated virus (CAV) or rAAV-retro, a rAAV-variant optimized for axonal uptake^8–10^.

While some of the mentioned methods are widely used, several caveats restrict their applicability. For instance, viruses that directly infect axonal terminals do not label cells that share a synaptic connection, but merely provide an area-specific retrograde label. Furthermore, viruses as well as trans-synaptic molecules can have cytotoxic effects (RV, HSV, VSV, WGA), their uptake can vary depending on cell type (CAV, HSV), expression strength and labelling intensity are rather low (certain RV-variants, TTC, WGA) or the synaptic transfer may not be limited to a single synapse, causing a ‘dilution’ effect and uncertainty about first order synaptic connections (TTC, WGA, HSV, VSV, non-pseudotyped RV)^6,7,11–15^.

To overcome these limitations, we combined the retrograde tracer (GFP-TTC^4^) with a presynaptic amplification mechanism for efficient detection and manipulation of connected cells. The amplification was achieved by expressing a flip-recombinase whose activity is dependent on the presence and binding of GFP (Flp-DOG^16^) in the acceptor cell. The activated Flp-DOG then drives the expression of fluorescent reporters, actuators, silencers, Cre-recombinase or any other protein of interest. Each of the genetic components is delivered to putatively connected brain regions through stereotactic injection of recombinant rAAVs (Fig. 1a, Supplementary Fig. 1), permitting long-term gene expression with no observable adverse effects on the rAAV infected cells.

**Fig. 1:**
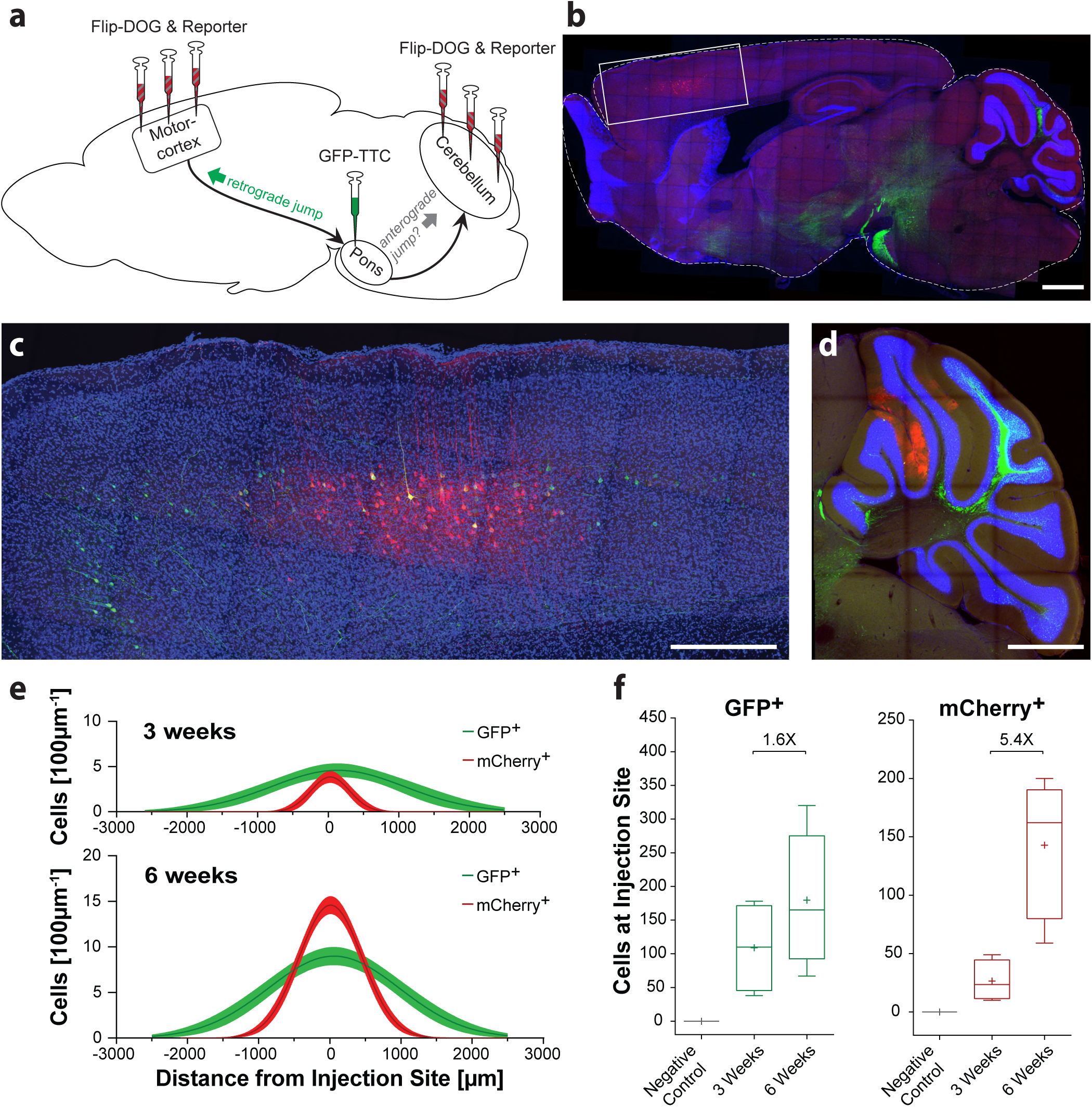
Design, proof of principle and quantification of trans-synaptic activation of gene expression in the cortico-pontine system. **a** Schematic overview of the rAAV injection sites. Directions of axonal projections are indicated by black arrows. Direction of retrograde GFP-TTC jump is indicated by a bold green arrow, while the putative anterograde jump is indicated by a bold grey arrow. **b** Low magnification overview image showing a sagittal section at the medio-lateral position of the premotor cortex injection 6 weeks post rAAV injection counterstained with DAPI (blue). Dashed lines denote the slice outline, the box shows the area depicted in high-magnification in **e**. Note that cerebellar and midbrain GFP-signal originates from axonal labelling (Supplementary Fig. S6). Scale bar: 1 mm. **c** High magnification image of the region marked in **b** showing both GFP-positive acceptor cells (green) as well as cells expressing mCherry (red) limited to layer 5 pyramidal cells. Scale bar: 500 µm. **d** High magnification overview image of a sagittal cerebellar section injected with the reporter mix showing both GFP-positive pontine projections and neurons of the deep cerebellar nuclei as well as mCherry positive (red) purkinje cells. Scale bar: 500 µm. **e** Distribution of GFP (green) and mCherry positive (red) cells along the cortical anterior-posterior axis relative to the injection site of the reporter mix 3 and 6 weeks post injection. Lines denote average (3 weeks n = 4; 6 weeks n = 6). Shaded areas denote SEM. **f** Total number of GFP (green) and mCherry positive (red) cells in the slice containing the injection site of the reporter mix 3 and 6 weeks post injection compared to the negative control (grey) after 6 weeks of expression (3 weeks n = 4; 6 weeks n = 6; Negative Control n = 3). Whiskers denote min and max values, line denotes median, cross denotes the mean.

To test retroLEAP, we used the well-described cortico-pontine system. In this circuit, neurons from premotor cortex layer 5 send long-range axonal projections to the pontine nuclei of the brain stem which in turn send projections to the contralateral cerebellum, allowing us to probe both retrograde and anterograde labelling ^17,18^ (Fig. 1a).

For probing the retrograde directionality, we injected rAAVs encoding the retrograde tracer GFP-TTC into the pontine nuclei, receiving layer 5 inputs, and a mixture of two rAAVs encoding Flp-DOG and the flip-dependent reporter FRT-ChR2-mCherry (henceforth referred to as the reporter mix), respectively, into the ipsilateral premotor cortex projecting to Pons (Fig. 1a). Six weeks after injection, the donor cells in the pontine nuclei and their cerebellar projections strongly expressed GFP-TTC (Fig. 1b,d, Supplementary Fig. 2). GFP-TTC was detected by Flp-DOG and activated dense mCherry-ChR2 expression in a large number of layer 5 neurons in the premotor cortical area (Fig. 1c, red). While the fluorescence intensity and number of both donor and acceptor cells increased between 3 and 6 weeks post injection (Fig. 1e, f, Supplementary Fig. 2), acceptor cells showing Flp-DOG-dependent mCherry expression (Fig. 1c, Supplementary Fig. 3) were confined to layer 5 pyramidal cells in the premotor cortex area that had been injected with the reporter mix (Fig. 1b, c). Thus, there is no evidence for second order transfer to neurons locally connected with the acceptor cells. If no GFP-TTC was injected into the pontine nuclei, no mCherry-positive cells could be detected in the premotor cortex, indicating that the observed mCherry-signal was not due to leaky expression (Supplementary Fig. 3). In addition, some acceptor neurons could be identified directly via the retrograde transfer of GFP-TTC (Fig. 1c, green, Supplementary Fig. 3). These labeled neurons are not restricted to the region of interest defined by the reporter mix injection and illustrate that under certain conditions synaptically transferred GFP-TTC can be detected, as reported previously^4^. Noticeably, the GFP-signal in both acceptor and donor cells did not degrade during tissue clearing (Supplementary Fig. 4 & Supplementary Dataset 1), making retroLEAP suitable for the use of single plane illumination microscopy (SPIM)^11^.

**Fig. 2:**
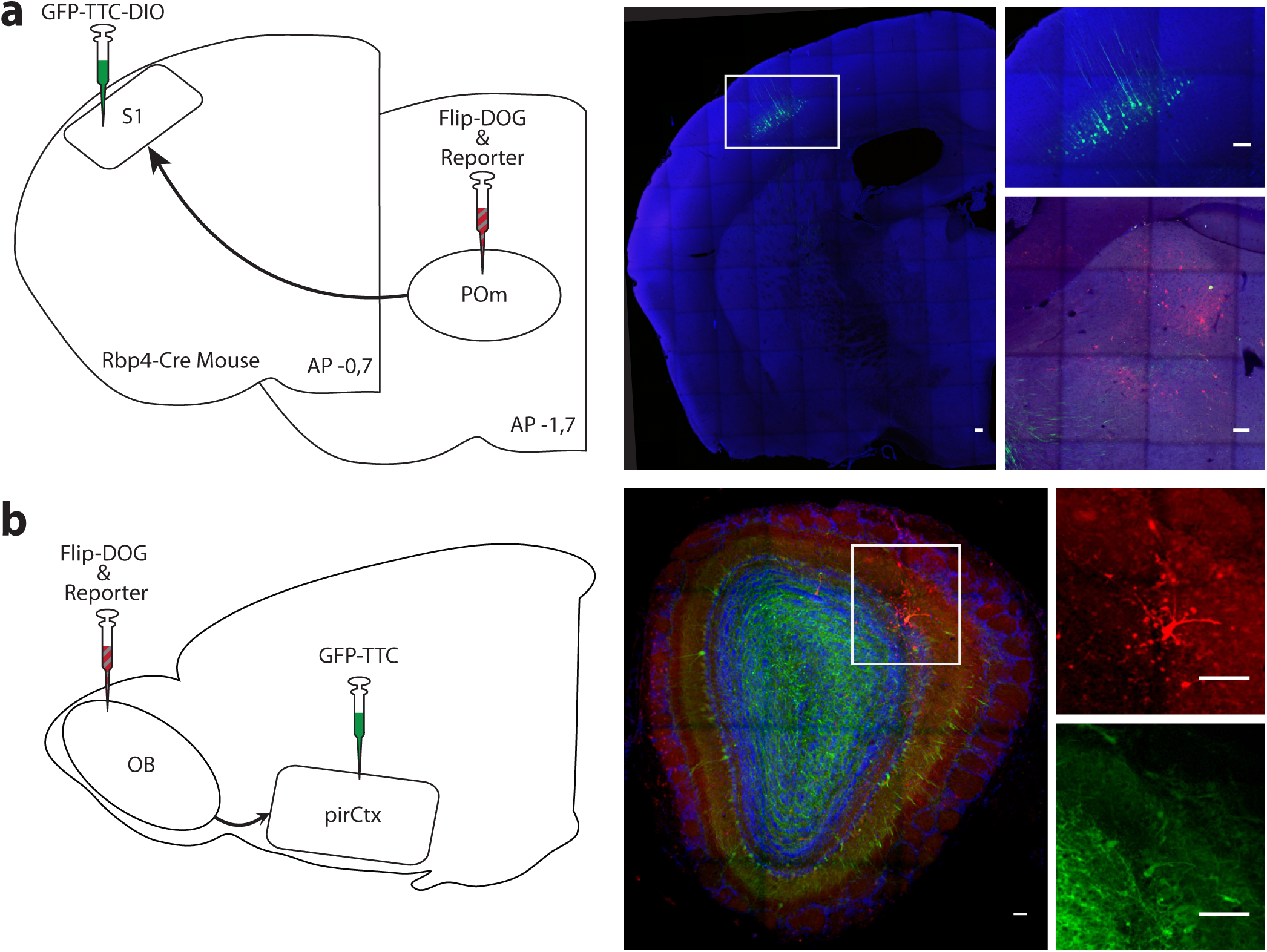
Application of retroLEAP in other brain circuits. **a** *Left:* Schematic overview of the injection sites of the rAAV encoding the Cre-dependent tracer in the somatosensory cortex (S1), the corresponding injection site of the reporter mix in the posteromedial nucleus of the thalamus (POm) and directionality of the projecting axons (black arrows). *Right:* Representative overview of the cortical tracer injection in S1 3 weeks post injection with the inset denoted by a white frame shown in a higher magnification with clearly visible GFP-TTC expressing layer 5 cells. The corresponding thalamic injection site shows both GFP-positive projections (green) as well as mCherry-positive (red) POm acceptor cells which lack visible GFP-signal. Nuclei are counter stained by DAPI (blue). Scale bars: 100 µm. **b** *Left:* Schematic overview of the injection sites in the olfactory bulb (OB) and piriform cortex (pirCtx) as well as the directionality of the projecting axons (black arrows). *Right:* Representative image of a coronal OB slice 3 weeks post injection showing GFP-TTC (green), mCherry (red) as well as DAPI-stained nuclei (blue). Note the abundance of GFP-positive cortigofugal projections throughout the entire granule cell layer (GCL) underneath the mitral cell layer (MCL). Inset denoted by a white frame shows an individual mitral cell labelled by both mCherry (red) as well as trans-synaptically transported GFP-TTC (green) in higher magnification. Scale bars: 100 µm.

More detailed quantification of the number of labelled cells in layer 5 shows that doubling the expression time from 3 to 6 weeks increased the number of GFP-positive acceptor cells by 1.6 fold (Fig. 1f). In contrast, the number of mCherry-positive cells increased by more than 5 fold, indicating that both the GFP-TTC transfer from donor to acceptor cells as well as the exclusively pre-synaptic GFP-mediated amplification system remain operative in virus-transduced cells many weeks after injection (Fig. 1f). It also highlights that the combination with Flp-DOG amplifies the signal in acceptor cells, hence allowing the identification of many more synaptic partners than sole GFP-TTC transfer.

To investigate the possibility of anterograde trans-synaptic GFP-TTC transfer, we injected the reporter mix into the cerebellum (Fig. 1a). As granule cells of the cerebellum receive inputs from neurons of the pontine nuclei via the mossy fibers, GFP or mCherry-positive cerebellar granule cells would indicate a putative anterograde transmission of GFP-TTC from the pons to the cerebellum^17,18^. However, three weeks after GFP-TTC delivery into the pons and despite the strong GFP-signal in the donor cells of the pontine nuclei and in their mossy fiber projections (Fig. 1b, d, Supplementary Fig. 5), we failed to identify mCherry or GFP-positive cerebellar granule cells (Supplementary Fig. 5f, j). On the other hand, we unexpectedly did observe GFP-positive cells within the deep cerebellar nuclei as well as GFP and mCherry-positive Purkinje cells which increased in number with longer expression time (Supplementary Fig. 5). As many targets of the deep cerebellar nuclei, like the red nucleus or the reticular formation, are located in close proximity to the injection site of the virus directing trans-synaptic tracer-expression in the pons, it is most likely that cells in these nuclei have been inadvertently co-infected with the GFP-TTC expressing virus, a problem that can be addressed by more limited local injections or by using cell-type specific expression of GFP-TTC.

To test the feasibility of cell-type specific tracer expression, we used the specific expression of Cre in pyramidal neurons of layer 5 in the somatosensory cortex of RBP4-Cre mice^19^. Here we used a Cre-dependent, rAAV-encoded GFP-TTC (GFP-TTC-DIO) to trace the cells projecting from the posteromedial nucleus of the thalamus (POm) onto layer 5 cells in S1 cortex^20^. As expected, the Cre-induced GFP-TTC expression was restricted to the donor cells in layer 5 of S1 region in RBP4-Cre mice (Fig. 2a) and the POm injected reporter mix was activated in cell bodies in the POm by the retrograde transport of GFP-TTC as indicated by the mCherry signal. In contrast to the cortico-pontine system, acceptor cells in the POm could not be visualized directly through the uptake of GFP-TTC, highlighting the necessity of a pre-synaptic amplification mechanism (Fig. 2a) for synaptic connections that apparently transfer GFP-TTC in a less efficient manner.

Given the high variability of labelling efficiency through direct GFP-TTC transmission, we chose the mitral cell (MC) to piriform cortex (pirCtx) connection (Fig. 2b), as uniform labelling of MCs has so far been difficult to achieve. In this circuit, the retrograde transport of GFP-TTC from pirCtx neurons to MCs was very effective, illustrated by the strong and early labelling of acceptor cells in the entire mitral cell layer (MCL) (Fig. 2b) which could be detected already one week post injection (Supplementary Fig. 6). Yet, due to the low spread of the reporter mix in the MCL, we observed only a few mCherry-positive MCs (Fig. 2b).

In summary, we present a novel combination of two independent systems that allow rAAV-based, cell type-specific, efficient and non-toxic retrograde tracing. This strategy encompasses several attractive features: (1) lack of toxicity, (2) high efficiency in labelling synaptically connected neurons, (3) extensive versatility in controlling gene expression in the presynaptically connected neurons (e.g. ChR, Cre recombinase, toxins, etc.) and (4) identification of weakly connected acceptor cells in which the levels of transmitted GFP-TTC are too low to be visualized. Thus, as our approach combines the low immunogenicity of rAAVs with the plethora of already existing rAAV-compatible sensors and actuators, it has the potential to become a key technology in the study and manipulation of neuronal networks.

## Supporting information

Complete Supplement incl. Figures

## Author Contributions

JKR, IS and TK designed the study and wrote the manuscript. JKR, IS & PP performed and analysed preliminary experiments. JKR & IS conducted the final experiments and analysed the data. IS wrote custom Matlab analysis scripts. HS & RS prepared high-titer rAAVs. SL performed tissue clearing. MK designed and executed subcloning of plasmid constructs. All authors read and approved of the final manuscript.

## Acknowledgments

We thank Ursula Lindenberger for excellent administrative assistance as well as Annette Herold and Claudia Koksch for technical assistance for virus production. Further we would like to thank Martin Wlotzka for invaluable IT support and Claudio Acuna for critical comments on an earlier version of the manuscript. Lastly we would like to thank Rohini Kuner for sharing the Rbp4-Cre line, Jonathan Tang for providing early Flp-DOG samples as well as Connie Cepko, Thomas Binz and Matthias Klugmann for providing plasmid samples used for subcloning.

## Data Availability Statement

Datasets and custom written code are available on request.

