## Supplementary material for "retroLEAP: rAAV-based retrograde trans-synaptic labeling, expression and perturbation": Complete Supplement incl. Figures

for

*** Correspondence:**

### Figure Legends

**Supplementary Fig. 1:** Components required for retrograde labelling. **a** The flp-recombinase (grey) is only active in the presence of GFP-TTC (green & white). This property is conferred by fusion of a destabilizing nanobody (blue) to the recombinase[1]. **b** Schematic overview of the three AAV constructs necessary for trans-synaptic activation of gene expression: each component of the system is delivered to acceptor cells and donor cells individually through stereotaxic injection of rAAVs.

**Supplementary Fig. 2:** Comparison of the pontine nuclei injected with rAAV-GFP-TTC in negative controls (without rAAV injection), 3 and 6 weeks after virus injection. **a,d,g** Representative DAPI staining for the negative control (**a**) as well as the GFP-TTC injected animals 3 weeks (**d**) and 6 weeks (**g**) after injection of the virus. **b,e,h** Representative images of the GFP signal observed in the negative control after 6 weeks (**b**) and the GFP-TTC injected animals 3 weeks (**e**) and 6 weeks (**h**) after injection of the virus. **c,f,i** Merged images of the individual channels per experimental group showing the DAPI (blue) and GFP (green) signal. Scale bar: 500 µm.

**Note:** All images were acquired using the same imaging settings optimized for 6 weeks after injection of the virus resulting in only very faint signal being visible 3 weeks after injection of the virus. The histograms were not adjusted to highlight the increase in fluorescence intensity with prolonged expression time.

**Supplementary Fig. 3:** Comparison of the premotor cortex area injected with the reporter mix in negative controls (without pontine rAAV-GFP-TTC injection) and in rAAV-GFP-TTC injected animals 3 and 6 weeks after injection of the virus. **a,e,i** Representative DAPI staining for the negative control (**a**) as well as the GFP-TTC injected animals 3 weeks (**e**) and 6 weeks (**i**) after injection of the virus. **b,f,j** Representative images of the GFP signal observed in the negative control after 6 weeks (**b**) and the GFP-TTC injected animals 3 weeks (**f**) and 6 weeks (**j**) after injection of the virus. **c,g,k** Representative images of the mCherry signal observed in the negative control after 6 weeks (**c**) and the GFP-TTC injected animals 3 weeks (**g**) and 6 weeks (**k**) after injection of the virus. **d,h,l** Merged images of the individual channels showing the DAPI (blue), GFP (green) and mCherry (red) signal. Scale bar: 500 µm.

**Note:** All images were acquired using the same imaging settings and histograms were adjusted for display purposes.

**Supplementary Fig. 4:** GFP signal 3 weeks after injection of the virus visualized using single plane illumination microscopy (SPIM). Selective optical sections at medio-lateral distances of the pontine nucleus injection site at 0.25mm (**a**) the premotor cortex injection site at 0.90mm (**b**) and 1.4mm (**c**) from the midline show widespread GFP-TTC positive cells throughout the cortex as well as projections into the cerebellum originating from the pontine nuclei.

**Note**: The histogram of the SPIM dataset was adjusted to visualize GFP-TTC positive cells in the cortex resulting in background fluorescence being visible as well as the injection site (**a**) appearing oversaturated.

**Supplementary Fig. 5:** Probing for anterograde trans-synaptic transfer of GFP-TTC to the cerebellum. **a, e,i** Representative DAPI staining for the negative control after 6 weeks (**a**) as well as the reporter mix injected animals 3 weeks (**e**) and 6 weeks (**i**) after injection of the virus. **b,f,j** Representative images of the GFP signal observed in the negative control after 6 weeks (**b**) and the reporter mix injected animals 3 weeks (**f**) and 6 weeks (**j**) after injection of the virus. White arrowheads (in **f** and **j**) denote GFP-positive neurons in the deep cerebellar nuclei. Areas denoted by dashed line are shown in higher magnification in the upper right-hand corners indicating mossy fiber endings.

**c,g,k** Representative images of the mCherry signal observed in the negative control after 6 weeks (**c**) and the reporter mix injected animals 3 weeks (**g**) and 6 weeks (**k**) after injection of the virus. Areas denoted by dashed line are shown in higher magnification in the upper right-hand corners to show mCherry-positive purkinje cells. **d,h,l** Merged images of the individual channels showing the DAPI (blue), GFP (green) and mCherry (red) signal. White arrowheads (in **h** and **l**) denote GFP-positive neurons in the deep cerebellar nuclei as highlighted in **f** and **j**. Scalebar: 100 µm.

**Supplementary Fig. 6:** Comparison of the transfer of GFP-TTC in the olfactory system at different incubation times **a,d,g** Representative DAPI staining 1 week (**a**) after injection of the virus in the piriform cortex as well as 3 weeks (**d**) and 6 weeks (**g**) after injection of the virus. **b,e,h** Representative images of the GFP signal observed 1 week (**b**) after injection of the virus in the piriform cortex as well as 3 weeks (**e**) and 6 weeks (**h**) after injection of the virus. Areas denoted by dashed line are shown in higher magnification in the upper right hand corner showing GFP-TTC labelled mitral cells. **c,f,i** Merged images of the individual channels showing the DAPI (blue) and GFP (green) signal. Scalebar: 100µm.

**Note:** Imaging parameters were optimized for the signal observed 3 weeks after injection of the virus and used for imaging of the experiments with other incubation periods. Therefore samples taken 1 week after injection of the virus show faint signals while samples taken 6 weeks after injection of the virus appear oversaturated. In the overview images no adjustment of the histograms was performed to better visualize the increase in transferred GFP-TTC signal over time.

### Supplementary Materials and Methods

#### Animals

All animal experiments were conducted in accordance with the European Union guidelines for the care and use of laboratory animals and approved by the responsible local authority (Regierungspräsidium Karlsruhe, Karlsruhe, Germany; HD35-9185.81/G-260/16).

Male mice aged 8 weeks upon injection were used. For the thalamocortical loop, Rbp4-Cre (RRID:MMRRC_031125-UCD) mice were used, expressing the Cre-recombinase in L5 of the somatosensory cortex[2]. For all other injections wild-type C57BL/6N animals were used.

#### Recombinant rAAV vectors

The expression cassettes (except for the Cre-dependent GFP-TTC-DIO casette) were subcloned into the multiple cloning site (MCS) of an rAAV-expression vector consisting of rAAV2 inverted tandem repeats (ITR), the woodchuck hepatitis virus posttranscriptional regulatory element (WPRE) as well as the bovine growth hormone polyadenylation sequence (BGHpA) (pAM-ITR-MCS-WPRE-BGHpA-ITR; a kind gift from Matthias Klugmann[3]).

For the Cre-dependent cassette, a modified rAAV-expression vector was used in which the MCS was flanked by loxP sites (pAM-ITR-loxP-MCS-loxP-WPRE-BGHpA-ITR).

The Flip-dependent reporter was already provided in a rAAV-compatible expression cassette.

Expression of the GFP-TTC tracer was limited to neurons by using the promoter of the human synapsin 1 gene (hSyn)[4]. For the remaining constructs, expression was controlled by the so-called CAG promoter to ensure unbiased and highly efficient gene expression regardless of cell-type[5].

**Table 1: rAAV vectors with corresponding restriction sites and expression vectors**

| Expression cassette | AAV-expression vector | Restriction sites | Addgene ID | Virus titer [genomes/µl] |
|---|---|---|---|---|
| Syn-GFP-TTC | pAM-ITR-MCS-WPRE-BGHpA-ITR | 3': MluI<br>5': HindIII | ***tba*** | 4.10E10 |
| CAG-GFP-TTC-DIO | pAM-ITR-loxP-MCS-loxP-WPRE-BGHpA-ITR | 3': EcoRV<br>5': XhoI | ***tba*** | 1.65E10 |
| CAG-Flp-DOG | pAM-ITR-MCS-WPRE-BGHpA-ITR | 3': EcoRI<br>5': NotI | ***tba*** | 1.39E8 |
| CAG-FLEX$^{FRT}$-ChR2(H134R)-mCherry | *unmodified* | *not applicable* | 75470 | 1.65E9 |

Flip-DOG (Addgene, #75469) and the Flip-dependent reporter (Addgene, #75470) were a kind gift from Connie Cepko[1]. TTC (Tetanus-Toxin C-Fragment) was a kind gift from Thomas Binz while the GFP-plasmid and the CAG promoter were a kind gift from Thomas Klugmann.

Recombinant AVVs (rAAVs) of the mixed serotype 1/2 were produced as described previously[6] and purified using FPLC as described previously[7].

#### Stereotaxic Injections

Injections were performed as described previously[6,8]. In brief, mice were anesthetized by i.p. injection of Medetomidine (0.715mg per kg body weight), Midazolam (9.3 mg per kg body weight) and Fentanyl

(0.024mg per kg body weight) and fixed in a stereotaxic holder (Kopf Instruments, Tujunga, CA, USA). Analgesia was provided by s.c. injection of Carprofen (5 mg per kg body weight).

Injection coordinates were obtained using the Allen Brain Atlas[9] and verified using retrobead (Lumaflour, #R180 and #G180) injections. The following coordinates relative to Bregma and the pial surface (in mm):

**Table 2: Injection Coordinates**

| Location | AP [mm] | ML [mm] | DV [mm] | Volume per Depth [nl] |
|---|---|---|---|---|
| Motor Cortex | 2.1<br>1.7<br>1.3 | 0.9<br>0.9<br>0.9 | 0.4 + 0.8<br>0.4 + 0.8<br>0.4 + 0.8 | 100<br>100<br>100 |
| Pons | - 3.8 | 0.25 | 4.5 | 200 |
| Cerebellum | - 6.2<br>- 6.2 | - 2.5<br>- 1 | 0.6 + 1.0 + 1.4<br>0.6 + 1.0 + 1.4 | 100<br>100 |
| Piriform Cortex | 1.94<br>1.94 | 2.1<br>2.1 | 3.25 + 3.5 + 3.75<br>3.25 + 3.5 + 3.75 | 100 |
| Somatosensory Cortex | - 0.7 | 3 | 0.7 + 1 | 150 |
| Thalamus | - 1.5<br>- 1.9 | 1.4<br>1.8 | 3<br>3 | 100<br>100 |
| Olfactory Bulb* | 0.1<br>0.3 | 0<br>0 | 0.05 + 0.1 + 0.15 + 0.2 + 0.25 + 0.3 + 0.35<br>0.05 + 0.1 + 0.15 + 0.2 + 0.25 + 0.3 + 0.35 | 25<br>25 |

| | | | | |
|---|---|---|---|---|
| | 0.5 | 0 | 0.05 + 0.1 + 0.15 + 0.2 + 0.25 + 0.3 + 0.35 | 25 |

*For the olfactory bulb (OB) injections, the injections coordinates are relative to the centre of the OB and the pial surface as described previously[9].

After surgery, anaesthesia was antagonized via i.p. injection of Atipamezole (1.898 mg per kg body weight), Flumazenil (0.506 mg per kg body weight) and Naloxon (0.304 mg per kg body weight). Once recovered, the animals were returned to their respective home cages.

**Slicing & Staining**

After the respective expression times, animals were transcardially perfused with PBS followed by PFA at room temperature, the brains removed and post-fixed in PFA (4% in PBS) at least overnight at 4°C. Sagittal slices of 100 µm thickness were obtained using a vibratome (Leica Microsystems, #VT100S) and stained in DAPI (0.00025 mg per ml in PBS) at least overnight at 4°C.

No immunohistochemistry was performed to amplify the native fluorescence signals of the two fluorescent proteins.

**Imaging**

Slices were pre-screened using an epifluorescence microscope (Leica Microsystems, #DM6000) equipped with a 1.25X/0.04NA objective (Leica Microsystems, #506215) and a 10X/0.4 NA objective (Leica Microsystems, #506284). Pre-screened premotor cortex and cerebellar regions containing cells expressing the reporter fluorophore were imaged in more detail using a confocal microscope (Leica Microsystems, #SP8) equipped with a 20X/0.75NA objective (Leica Microsystems, #11506343) and a 10X/0.4NA objective (Leica Microsystems, #11506293) at 2.5 µm Z-resolution and 200Hz bidirectional scan speed. DAPI and GFP signals were acquired using PMT detectors, mCherry signals were acquired using HyD detectors (Leica Microsystems, HyD©). For all images used for quantification, identical imaging parameters (i.e. laser intensities and PMT limits) were used.

### Quantification

For quantification of the cell density and amount of cells per slice, a custom-written Matlab routines (Mathworks, RRID:SCR_001622) based on Matlab version 2016b and Fiji Version 1.52a (Fiji, RRID:SCR_002285) were developed. Briefly, confocal stacks were converted into maximal intensity projections and cells were manually labelled. Layer 5 was manually plotted as a straight line and subsequently used as a reference to automatically determine the distribution of cells along the anterio-posterior axis of cortical layer 5 as well as the total number of cells in the slice containing the reporter mix injection site.

The slice containing the reporter mix injection site was defined as the slice containing the highest number of reporter fluorophore expressing cells. This slice was subsequently analysed in more detail by fitting the distribution of cells along the anterior-posterior cortical axis (binned in 100 µm increments) to a Gaussian distribution. For all animals of a respective experimental group, these Gaussian fits were aligned and averaged to generate an average distribution of cells per experimental group. The same slice was used for the analysis of GFP-positive cells using the same procedure as described for mCherry-positive cells.

As the negative control samples were found to not contain any mCherry-positive cells, the slice containing the reporter injection site was determined using the Allen Reference Atlas[10].

All images were coded and the quantifier remained blind to the identity of the images until quantification for all datasets was finalized.

### Tissue Clearing

Dehydration and optical clearing of PFA-fixed samples was performed using BABB-based clearing as described previously[11]. Briefly, whole mouse brains were dehydrated by incubation in water/tert-butanol mixtures of increasing tert-butanol concentration (30%, 50%, 70%, 80%, 96% and 100%) at room-temperature for at least 24 hrs per concentration. Subsequently, clearing was performed by incubation BABB clearing solution (1:2 mixture of benzyl alcohol [Sigma, #24122] and benzyl benzoate [Sigma, #B6630]) twice for 24 hrs at room temperature. For each solution, the pH was adjusted to ≥ 9.5 using triethylamine (Sigma, #90340) and a dedicated “InLab Science” pH sensor optimized for organic solvents (Mettler Toledo, #6622837).

#### Light Sheet Imaging and Image Processing

Light Sheet Imaging and Image Processing was performed using a LaVision Ultramicroscope II SPIM microscope (LaVision BioTec, Germany) equipped with a 4X/0.3 NA objective (5.6mm working distance) (LaVision BioTec, #205852). Imaging was performed in a dedicated 100% quartz imaging chamber (LaVision BioTec, #207467) filled with the same BABB-solution used for clearing to avoid artefacts due to differing refractive indices or pH levels. Samples were illuminated alternatively from both sides by the laser light. Samples were imaged using 5.5 Megapixel Andor Neo sCMOS camera (LaVision BioTec, #200023) at a fixed Z-step size of 5 µm and a xy-resolution of 1.625 µm/px. Image stacks were automatically merged using TeraStitcher, incorporated in the LaVision microscopy software (ImSpectorPro), allowing for parallel stitching of the data. The resulting tile-scans were downsampled by a factor of 10 for further processing.

#### Statistical Analysis

Data was analysed using custom-written Matlab routines as well as Prism Version 6.0 (Graphpad Software, RRID:SCR_002798). Statistical significance of was assessed using parametric unpaired two-tailed t-tests with Welch’s correction (* $p < 0.05$, ** $p < 0.01$, *** $p < 0.001$). Unless otherwise stated, data is presented as mean ± SEM.

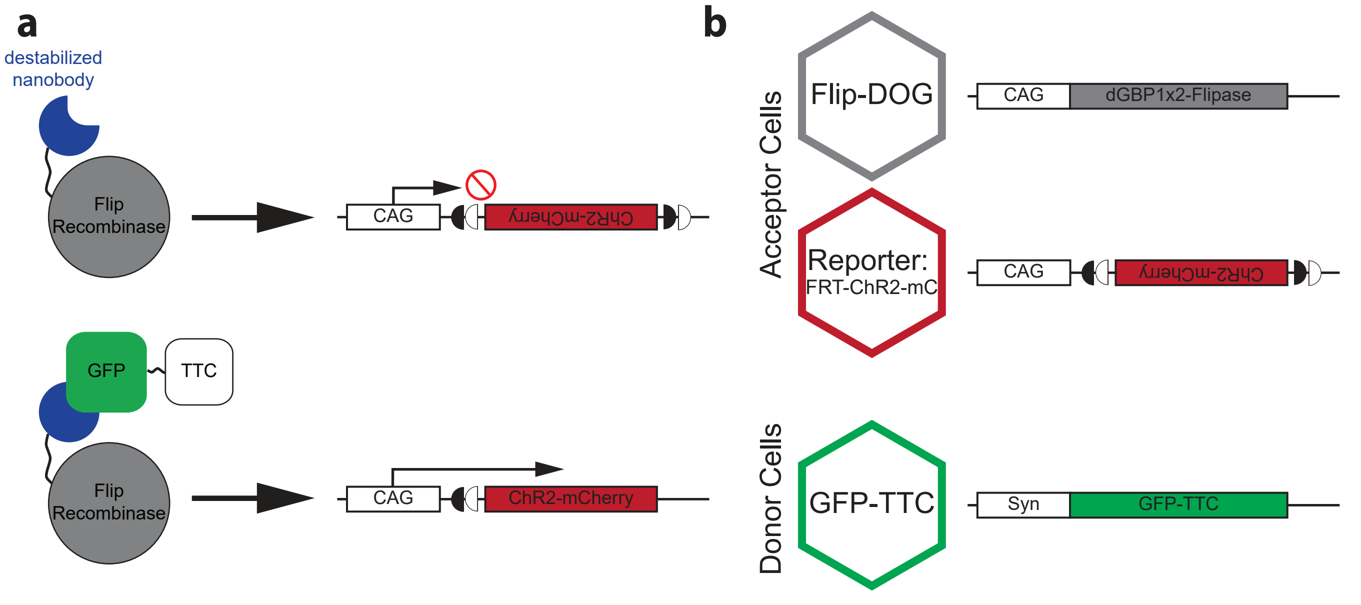
a
destabilized nanobody
Flip Recombinase
CAG
ChR2-mCherry
GFP
TTC
Flip Recombinase
CAG
ChR2-mCherry
b
Acceptor Cells
Flip-DOG
CAG
dGBP1x2-Flipase
Reporter:
FRT-ChR2-mC
CAG
ChR2-mCherry
Donor Cells
GFP-TTC
Syn
GFP-TTC

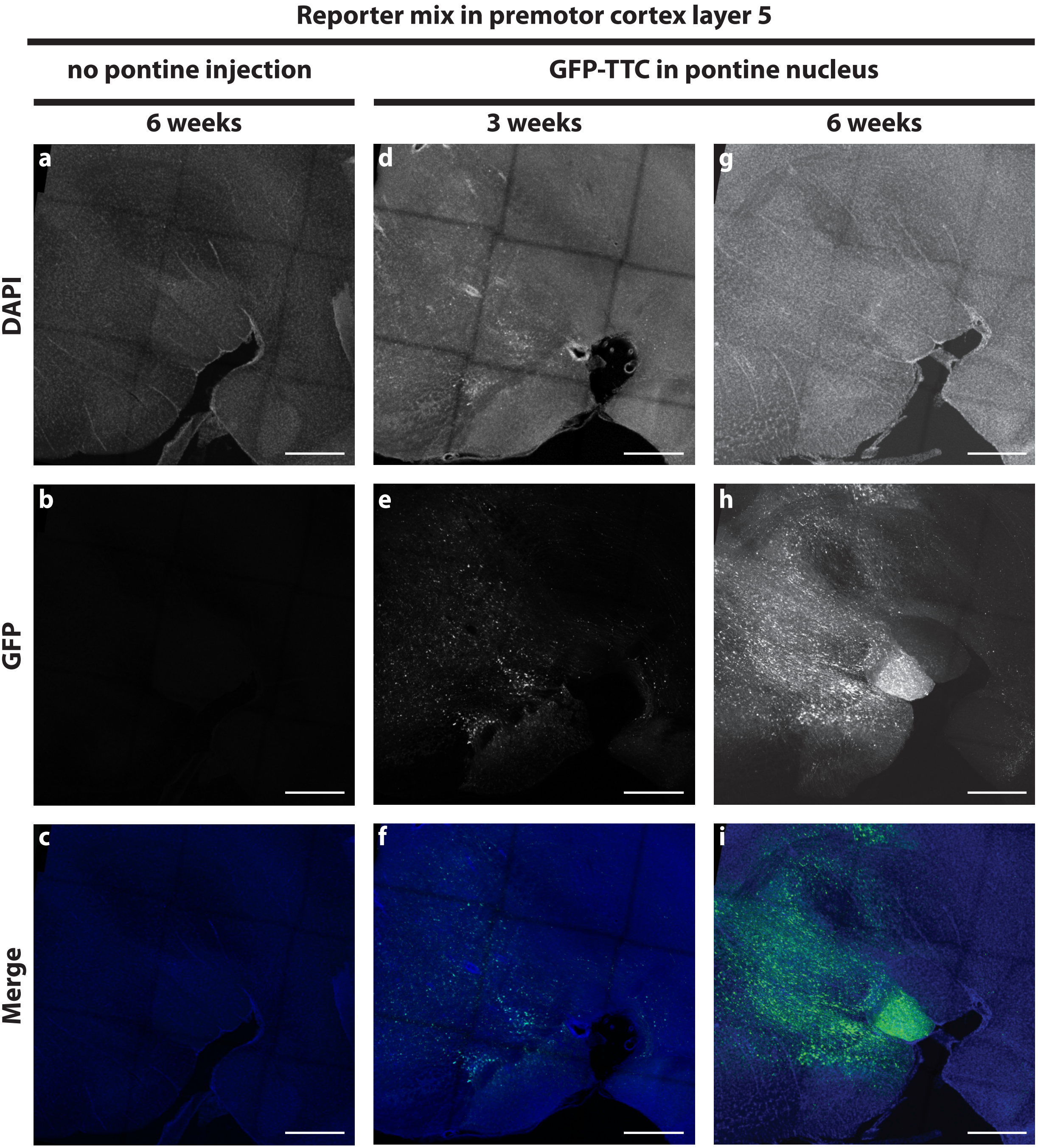
Reporter mix in premotor cortex layer 5
no pontine injection
GFP-TTC in pontine nucleus
6 weeks
3 weeks
6 weeks
DAPI
GFP
Merge
a
b
c
d
e
f
g
h
i

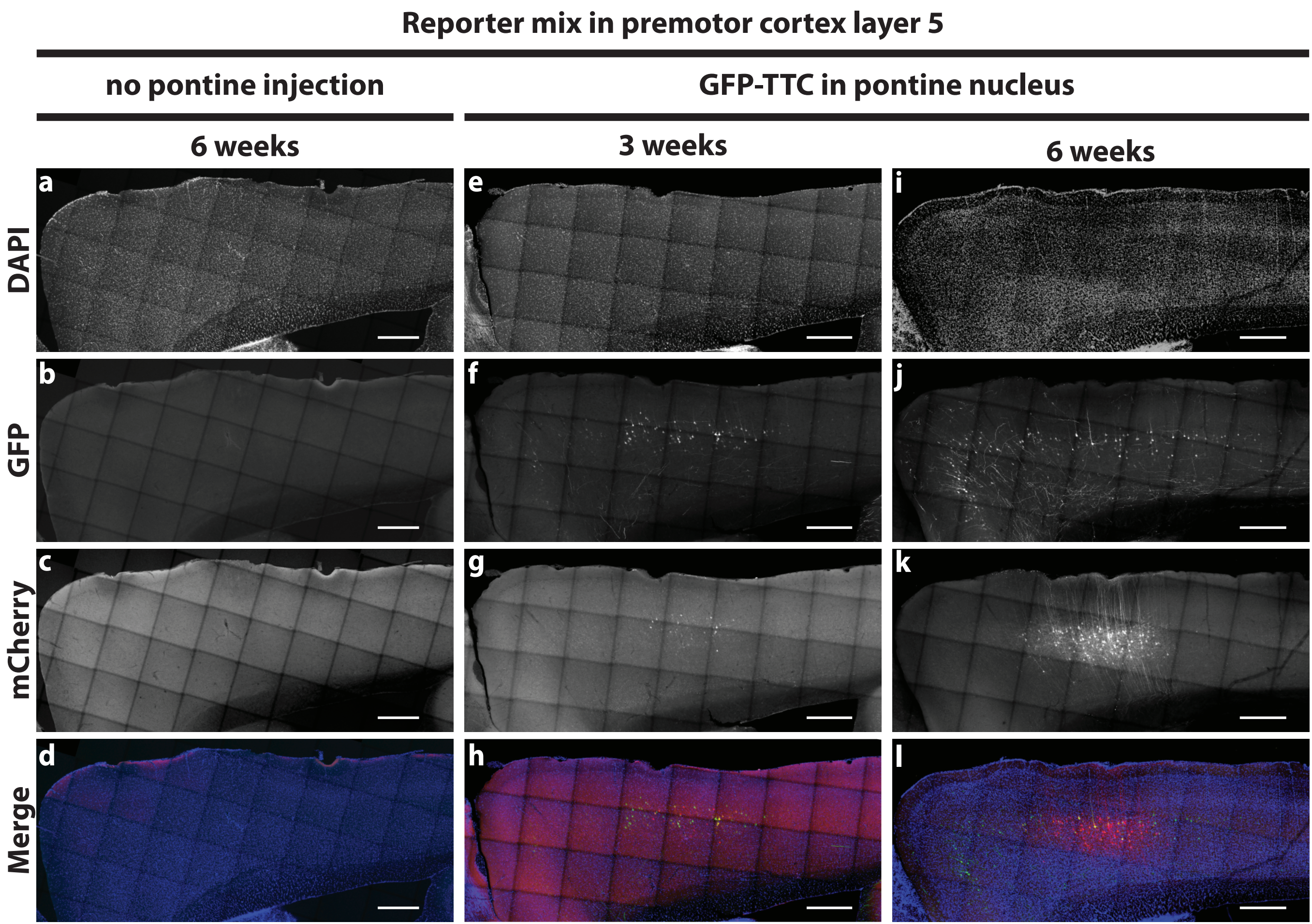

Reporter mix in premotor cortex layer 5
no pontine injection
GFP-TTC in pontine nucleus
6 weeks
3 weeks
6 weeks
DAPI
GFP
mCherry
Merge
a
b
c
d
e
f
g
h
i
j
k
l

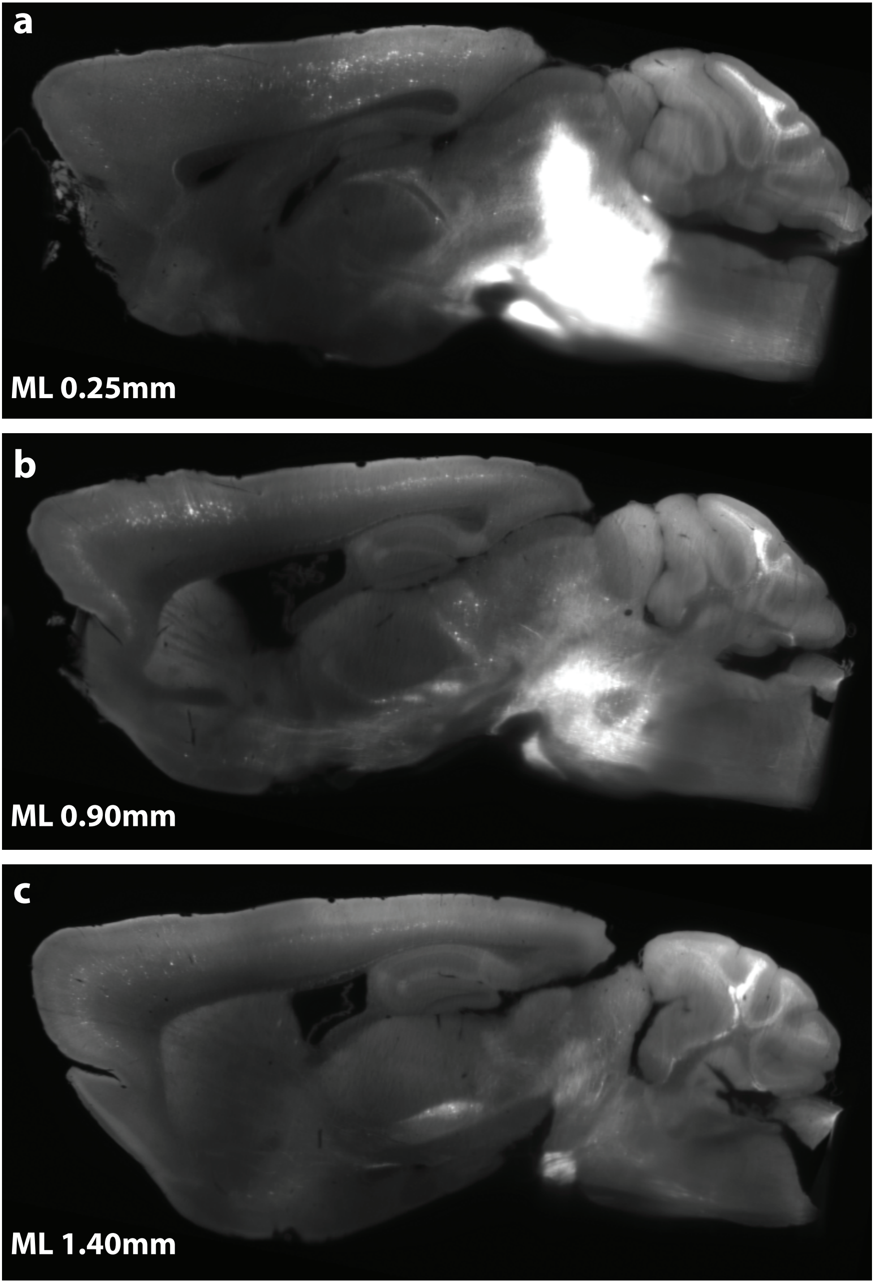
a
ML 0.25mm
b
ML 0.90mm
c
ML 1.40mm

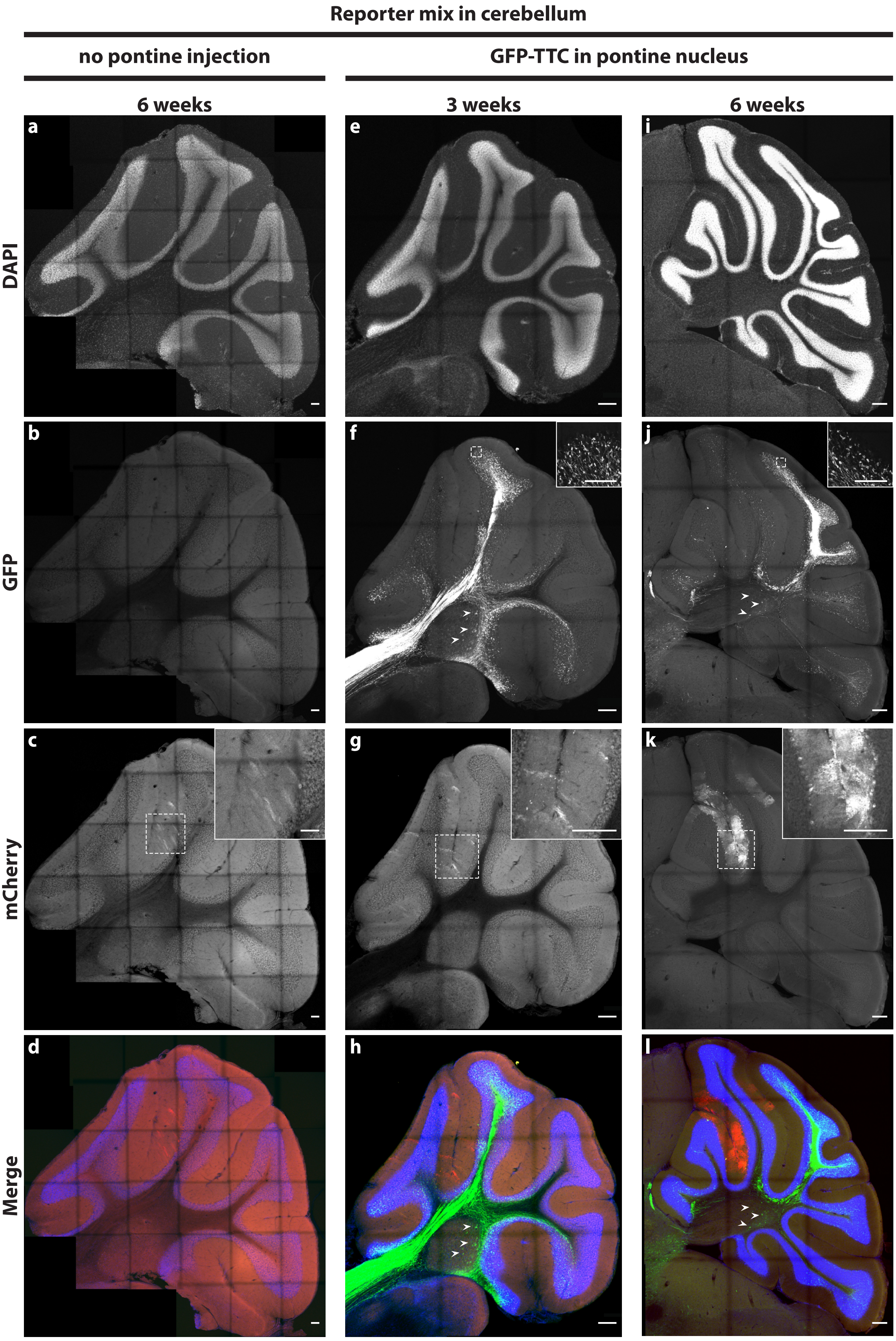

Reporter mix in cerebellum
no pontine injection
GFP-TTC in pontine nucleus
6 weeks
3 weeks
6 weeks
DAPI
GFP
mCherry
Merge
a
b
c
d
e
f
g
h
i
j
k
l

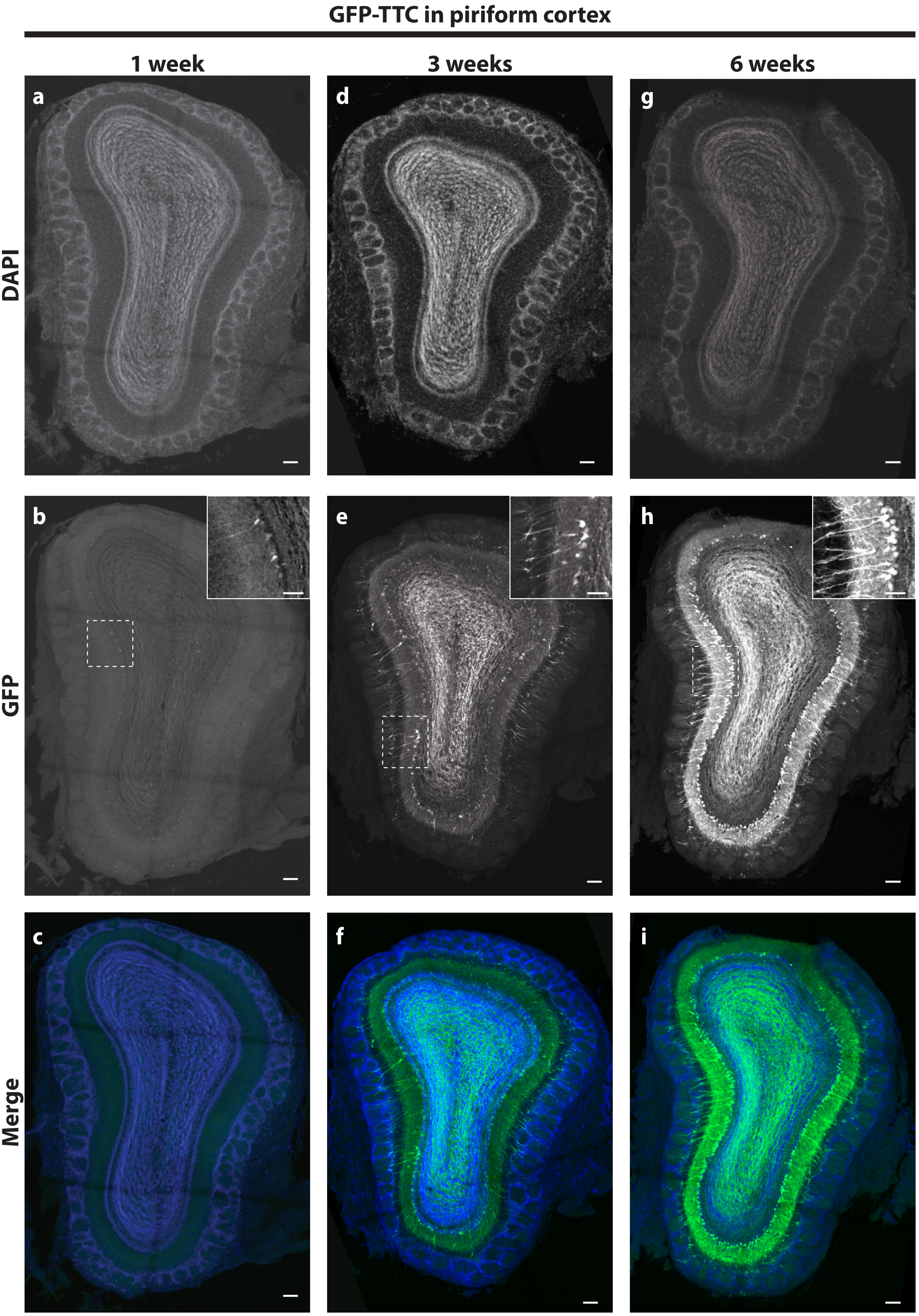
GFP-TTC in piriform cortex
1 week
3 weeks
6 weeks
DAPI
GFP
Merge
a
b
c
d
e
f
g
h
i
